# A chromosome-level genome resource for studying virulence mechanisms and evolution of the coffee rust pathogen *Hemileia vastatrix*

**DOI:** 10.1101/2022.07.29.502101

**Authors:** Peri A Tobias, Richard J. Edwards, Priyanka Surana, Hayley Mangelson, Vera Inácio, Maria do Céu Silva, Vitor Várzea, Robert F. Park, Dora Batista

## Abstract

Recurrent epidemics of coffee leaf rust, caused by the fungal pathogen *Hemileia vastatrix*, have constrained the sustainable production of Arabica coffee for over 150 years. The ability of *H. vastatrix* to overcome resistance in coffee cultivars and evolve new races is inexplicable for a -pathogen that supposedly only utilizes clonal reproduction. Understanding the evolutionary complexity between *H. vastatrix* and its only known host, including determining how the pathogen evolves virulence so rapidly is crucial for disease management. Achieving such goals relies on the availability of a comprehensive and high-quality genome reference assembly. To date, two reference genomes have been assembled and published for *H. vastatrix* that, while useful, remain fragmented and do not represent chromosomal scaffolds. Here, we present a complete scaffolded pseudochromosome-level genome resource for *H. vastatrix* strain 178a (Hv178a). Our initial assembly revealed an unusually high degree of gene duplication (over 50% BUSCO basidiomycota_odb10 genes). Upon inspection, this was predominantly due to a single scaffold that itself showed 91.9% BUSCO Completeness. Taxonomic analysis of predicted BUSCO genes placed this scaffold in Exobasidiomycetes and suggests it is a distinct genome, which we have named Hv178a <u>a</u>ssociated fungal <u>g</u>enome (Hv178a AFG). The high depth of coverage and close association with Hv178a raises the prospect of symbiosis, although we cannot completely rule out contamination at this time. The main Ca. 546 Mbp Hv178a genome was primarily (97.7%) localised to 11 pseudochromosomes (51.5 Mb N50), building the foundation for future advanced studies of genome structure and organization.

## Introduction

The specialized fungal pathogen *Hemileia vastatrix* Berk. & Broome (phylum Basidiomycota, class Pucciniomycetes, order Pucciniales), causing the devastating disease coffee leaf rust (CLR), has been the major constraint to Arabica coffee (*Coffea arabica*) production for more than one and a half centuries (McCook and Vandermeer 2015; Cabral *et al*. 2016; Talhinhas *et al*. 2017a). First recorded by an English explorer in 1861 near Lake Victoria (East Africa) on wild *Coffea* species, CLR wiped out Arabica coffee cultivation from Ceylon (Sri Lanka) only five years later, with devastating social and economic consequences (Morris 1880). Since this historical first outbreak, the disease gained a worldwide distribution, spreading progressively through coffee production areas of Asia and Africa, and, finally, crossing the Atlantic Ocean where it expanded across Latin America (McCook 2006). With the recent detection of CLR in Hawaii in 2020 (Keith *et al*. 2021), the disease became endemic in all coffee producing regions of the world. Although the development and cultivation of rust resistant varieties has successfully contributed to CLR control, high adaptation in the fungus shaped by the dynamic system of host-pathogen co-evolution has led to recurrent and severe epidemics resulting in heavy yield and revenue losses. Latest reports estimate production losses in excess of $1 billion annually worldwide (Kahn 2019). Furthermore, during the last decade, a cluster of epidemic outbreaks known as “the big rust” burst out across Latin America and the Caribbean on a level similar to the one observed in Ceylon; this extended both the temporal and spatial range of the disease, conferring to CLR the status of “natural disaster” in the tropics (Avelino *et al*. 2015; McCook and Vandermeer 2015; Silva *et al*. 2022).

The global dissemination of *H. vastatrix* across continents seems to be intimately linked to the historical evolution of the global coffee industry. Only recently, the worldwide genetic population structure of this pathogen was uncovered by genome-wide SNP data, which revealed strong local adaptation driven by coffee hosts (Rodrigues *et al*. 2022). As a result of the high ability of the pathogen to respond rapidly to selection pressure, more than 55 physiological races have been identified to date (Silva *et al*. 2022). However, no direct link between such high phenotypic diversity and molecular diversity has yet been found, and no virulence locus has characterized so far (Talhinhas *et al*. 2017b; Silva *et al*. 2022).

The rapid evolution of *H. vastatrix* to overcome resistance in coffee cultivars is even more puzzling considering that the pathogen exists almost exclusively in the asexual urediniospore stage of its life cycle and that the dikaryotic urediniospores are the only known functional propagules (Talhinhas *et al*. 2017b; Silva *et al*. 2018, 2022; Koutouleas *et al*. 2019a). *Hemileia vastatrix* is a hemicyclic fungus producing urediniospores, teliospores and basidiospores, but teliospores occur very rarely (Chinnappa and Sreenivasan 1965; Coutinho *et al*. 1995; Fernandes *et al*. 2009) and basidiospores do not re-infect coffee (Rodrigues *et al*. 1980; Kushalappa and Eskes 1989). This led some authors to hypothesize that *H. vastatrix* is a heteroecious rust (Gopalkrishnan 1951; Petersen 1974), although no alternate host was ever found despite numerous efforts (Koutouleas *et al*. 2019b). Given that *H. vastatrix* belongs to the earliest rust lineages within the Pucciniales (family Mikronegeriaceae) (Catherine Aime 2006; McTaggart *et al*. 2016; Aime and McTaggart 2021), it is possible that it has lost the ability to produce sexual spores during evolution because adaptation to survival would have favored the uredinial stage. Although life cycle reduction and predominance of the asexual stage may represent an evolutionary dead-end (Aime and McTaggart 2021), *H. vastatrix* must have found alternative strategies to generate genetic variability because evidence of meiosis within urediniospores (Carvalho *et al*. 2011) and signatures of recombination retrieved from DNA marker and genomic data (Maia *et al*. 2013; Cabral *et al*. 2016; Silva *et al*. 2018) has been reported, as well as introgression footprints (Silva *et al*. 2018).

Deconstructing the evolutionary complexity between *H. vastatrix* and its only known host and understanding how the pathogen evolves virulence rapidly are crucial to devise options for sustainable disease control. However, achieving such key goals relies on the availability of a comprehensive and high-quality genome reference assembly. Previous research efforts reported two *H. vastatrix* genome assemblies based on a combined sequencing strategy of 454 and Illumina and of PacBio RSII and Illumina, respectively (Cristancho *et al*. 2014; Porto *et al*. 2019). These genomes provided important new knowledge on the high content of repetitive elements and on candidate effector genes. Cristancho *et al*. (2014) generated a partial hybrid assembly of 333 Mbp comprising eight distinct *H. vastatrix* isolates occurring in Colombia. Although a significant advance, this resource was of limited assistance for genomic data analyses given the high proportion of repetitive content (ca. 75%) and the predicted genome size of ca. 800 Mbp (Tavares *et al*. 2014),. Later, Porto *et al*. (2019) produced an improved assembly of 547 Mbp with a contig N50 of ∼10 Kbp, containing about 82% repetitive elements, of a local Brazilian *H. vastatrix* isolate (race XXXIII). While this assembly is useful, it is highly fragmented, at 116,756 contigs. Here, we report a pseudochromosome-level genome assembly of *H. vastatrix* (strain CIFC Hv178a), including a complete circularised sequence of the mitochondrial genome, as a high-quality resource for advancing knowledge on the complex coffee rust pathogen. We include an unusual finding of an 18 Mbp fungal scaffold that appears to be associated with this strain. We make no conclusions about this but name it as an Hv178a <u>a</u>ssociated fungal <u>g</u>enome (Hv178a AFG) for future investigations.

### Chromosome-level *Hemileia vastatrix* (CIFC Hv178a) assembly

#### Fungal material, DNA extraction and sequencing

*Hemileia vastatrix* isolate CIFC Hv178a (type specimen of race XIV: genotype v_2,3,4,5_), maintained at the spore collection of Centro de Investigação das Ferrugens do Cafeeiro (CIFC), Instituto Superior de Agronomia, Universidade de Lisboa, was used in this work. This isolate is a mutant strain obtained in 1960 from CIFC’s greenhouse complex derived from isolate CIFC Hv178 (race VIII: genotype *v*_*2*_,_*3,5*_), collected in India (Balehonnur, Coffee Research Station) in 1958 from *Coffea arabica* S.288-23. CIFC Hv178a was multiplied on its differential host plant (*C. arabica* accession CIFC H147/1, carrying the resistance factors *S*_*H*_*2, S*_*H*_*3, S*_*H*_*4* and *S*_*H*_*5*), as previously described (d’Oliveira 1954). Single uredospore pustules were collected, frozen in liquid nitrogen, and stored at -80°C until further processing. High molecular weight (HMW) genomic DNA was extracted using a cetyltrimethylammonium bromide (CTAB)-based protocol modified from Schwessinger and Rathjen (2017). DNA samples were further purified using Pacific Biosciences (PacBio) SampleNet – Shared Protocol (https://www.pacb.com/wp-content/uploads/2015/09/SharedProtocol-Extracting-DNA-usinig-Phenol-Chloroform.pdf). DNA concentration and purity were measured using a Multiskan SkyHigh Microplate Spectrophotometer (Thermofisher Scientific) and a Qubit 4 Fluorometer (Thermofisher Scientific), and its quality checked on an agarose gel. Ten micrograms of high-quality HMW DNA were sent to CD Genomics (New York, USA) for SMRT bell size select long read library preparation and PacBio Sequel II sequencing (continuous long read runs). The same sample of HMW DNA was used for library construction with TruSeq DNA PCR-Free kit, following manufacturer’s recommendations, and resequencing with paired-end 151 bp reads at CD Genomics on Illumina an NovaSeq platform.

#### Draft genome assembly

We received raw PacBio Sequel (Menlo Park, California, USA) SMRT™ continuous long reads (CLR) from CD Genomics (New York, USA). Prior to correction we had 7,539,679 reads (166 Gigabases), with median read length of 17,992. This provided approximately 200 times coverage on an expected genome size of 800 Mb, based on flow cytometry (Tavares *et al*. 2014). An initial assembly using Canu version 1.9 (Koren *et al*. 2016) on the University of Sydney high performance compute cluster, Artemis, running PBS-Pro (v 13.1.0.160576) was run with the following parameters including default corOutCoverage; genomeSize=800m, correctedErrorRate=0.040, batOptions=“-dg 3 -db 3 -dr 1 -ca 500 -cp 50” and built a diploid assembly of 1.6 Gb with 7,374 contigs, an N50 of 62,8510 and L50 of 458. Following this we tested Canu version 2.1 using these parameters; genomeSize=800m, corOutCoverage=100, corMaxEvidenceErate=0.15, correctedErrorRate=0.040, batOptions=“-dg 3 -db 3 -dr 1 -ca 500 -cp 50”. These options produced an assembly of 1.71 Gb with 6,088 contigs, an N50 of 74,5640 and L50 of 438. A final assembly using Canu version 2.0 was run on Australia’s National Compute Infrastructure (NCI) using the following parameters to maximise coverage; genomeSize=1.6g corOutCoverage=1000 “batOptions=-dg 3 -db 3 -dr 1 -ca 500 -cp 50”. This step produced trimmed reads that were assembled with Canu version 2.1 on Artemis to produce an assembly of 1.9 Gb with 6,602 contigs, an N50 of 62,8510 and L50 of 458. After testing genome outputs for contiguity with Quast version 5.0.2 (Gurevich *et al*. 2013), and conserved single copy gene completeness with BUSCO version 3.0 (Simão *et al*. 2015) with the basidiomycota_odb9 database, we found optimal results from the high coverage (>200 X) final assembly. We used this output, named CR_2103 (<u>c</u>offee rust and date), going forward.

#### Genome deduplication

We ran Purge Haplotigs version 1.0 (Roach *et al*. 2018) to deduplicate the 1.9 Gb genome using these counts per read depth; -l 32 -m 115 -h 170 after mapping with Minimap2 version 2.3 (Li 2018). We used these purge parameters for the cut-offs; -I 250M -a 70, after testing percent cut-offs of 65, 75, 80 and 85 percent for expected genome size and BUSCO completeness. We produced two genome fasta outputs with the following BUSCO statistics; C:94.7%[S:59.5%,D:35.2%],F:2.2%,M:3.1% and C:82.4%[S:62.5%, D:19.9%],F:4.3%,M:13.3% for the primary (553M) and haplotig (813M) assemblies. The BUSCO statistics indicated difficulties in separating the haplotypes, with large numbers of duplicated conserved genes in each genome. We proceeded with scaffolding using Hi-C data, described below, with the most suitable genome based on expected size and completeness.

#### Cross-linking of spores and chromatin conformation capture (Hi-C) sequencing

To improve the scaffolding of our genome assembly we obtained chromatin conformation sequence data. About 400 mg of CIFC Hv178a urediniospores were crosslinked in a 1% formaldehyde solution for 20 min at room temperature. Crosslinking was quenched by adding glycine to a final concentration of 125 mM. The fixed tissue was then ground in liquid nitrogen and the stored frozen powder sent to Phase Genomics (Seattle, WA) for nuclei isolation, Hi-C library preparation and sequencing. Data was generated using a Phase Genomics (Seattle, WA) Proximo Hi-C 4.0 Kit, which is a commercially available version of the Hi-C protocol (Lieberman-Aiden *et al*. 2009). Following the manufacturer’s instructions for the kit, intact cells from two samples were crosslinked using a formaldehyde solution, digested using the DPNII restriction enzyme, and proximity ligated with biotinylated nucleotides to create chimeric molecules composed of fragments from different regions of the genome that were physically proximal in vivo. Continuing with the manufacturer’s protocol, molecules were pulled down with streptavidin beads and processed into an Illumina-compatible sequencing library. Sequencing was performed on an Illumina HiSeq 4000, generating a total of 31,666,891 PE150 read pairs.

#### Chromosome-scale scaffolding with Hi-C reads

The deduplicated primary assembly was scaffolded by Phase Genomics (Seattle, WA). Reads were aligned to the draft assembly (CR_2103_70) following the manufacturer’s recommendations (Phase Genomics 2019). Briefly, reads were aligned using BWA-MEM (Li and Durbin 2010) with the -5SP and -t 8 options specified, and all other options default. SAMBLASTER (Faust and Hall 2014) was used to flag PCR duplicates, which were later excluded from analysis. Alignments were then filtered with samtools (Li *et al*. 2009) using the -F 2304 filtering flag to remove non-primary and secondary alignments.

Phase Genomics’ Proximo Hi-C genome scaffolding platform was used to generate chromosome-scale scaffolds following the single-phase scaffolding procedure described in Bickhart *et al*. (2017). As in the LACHESIS method (Burton *et al*. 2013), this process computes a contact frequency matrix from the aligned Hi-C read pairs, normalized by the number of DPNII restriction sites (GATC) on each contig, and constructs scaffolds in such a way as to optimize expected contact frequency and other statistical patterns in Hi-C data. Approximately 20,000 separate Proximo runs were performed to optimize the number and construction of scaffolds in order to make them as concordant with the observed Hi-C data as possible. This process resulted in a set of 12 chromosome-scale scaffolds containing 543.9 Mbp of sequence (94.8% of the corrected assembly). Juicebox (Rao *et al*. 2014; Durand *et al*. 2016) was then used to correct scaffolding errors. The final pseudochromosome assembly included 12 scaffolds spanning 552.8 Mbp (96% of input) with a scaffold N50 of 49.4 Mbp.

#### Mitochondrial genome assembly

In order to build the complete mitochondrial organelle genome, we ran Novoplasty (Dierckxsens *et al*. 2017) version 4.3.1, using the PE Illumina resequencing reads and the *Hemileia vastatrix* cytochrome b (cytb) gene, (NCBI accession DQ209282.1) as the seed (Grasso *et al*. 2006). Config specifications included genome size range=100-200 kb, kmer size=33 and all other defaults. These settings produced a complete circularised mitochondrial sequence at ∼171 kb in size. NUMTFinder (Edwards *et al*. 2021) identified a 51,490 bp contig with 51,449 bp (99.9%) identity to a 51,476 bp chunk of the mtDNA genome. This contig was removed from the assembly.

#### Genome quality assessment

We estimated the functional completeness of the genome using BUSCO (Simão *et al*. 2015) version 5.1.2 in genome mode with the database, basidiomycota_odb10, and Metaeuk (Levy Karin *et al*. 2020) gene predictions. Basic genome statistics were determined with Quast (Gurevich *et al*. 2013) version 5.0.2. Kmer-based genome sequence completeness and accuracy was estimated with Merqury v1.3 (Rhie *et al*. 2020).

#### Taxonomic contamination assessment and filtering

BUSCO proteins were assigned to taxonomic groups using Taxolotl v0.1.2 (https://github.com/slimsuite/taxolotl). Additional protein predictions were made using GeMoMa v1.7.1 (Keilwagen *et al*. 2019) homology-based annotation based on Ensembl Release 50 reference genomes for available Basidiomycota species: *Cryptococcus neoformans, Microbotryum violaceum, Puccinia graminis, Puccinia striiformis, Puccinia triticin, Sporisorium reilianum*, and *Ustilago maydis*. Maximum intron size was set to 200 kb. Proteins were searched against NCBInr using MMSeqs2 v13-45111 (Mirdita *et al*. 2021) and Taxolotl output visualised with Pavian (Breitwieser and Salzberg 2020). The whole genome assembly protein homology, Figure 1A, shows divergence at Class level to Exobasidiomycetes and Pucciniomycetes. When we separate out the associated fungal genome the taxonomic homology is more concordant for two lineages, Figure 1B and C.

**Figure 1.**
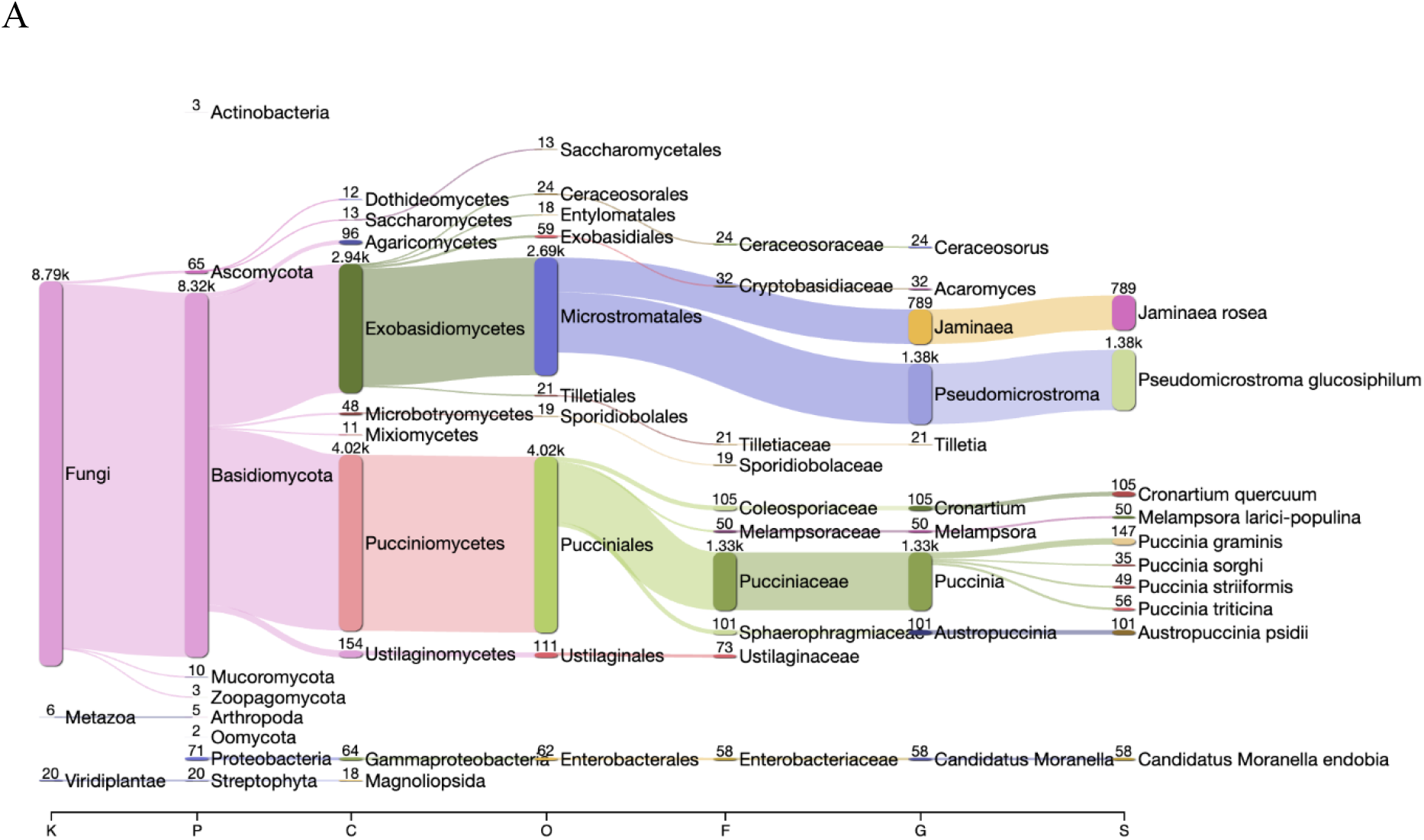
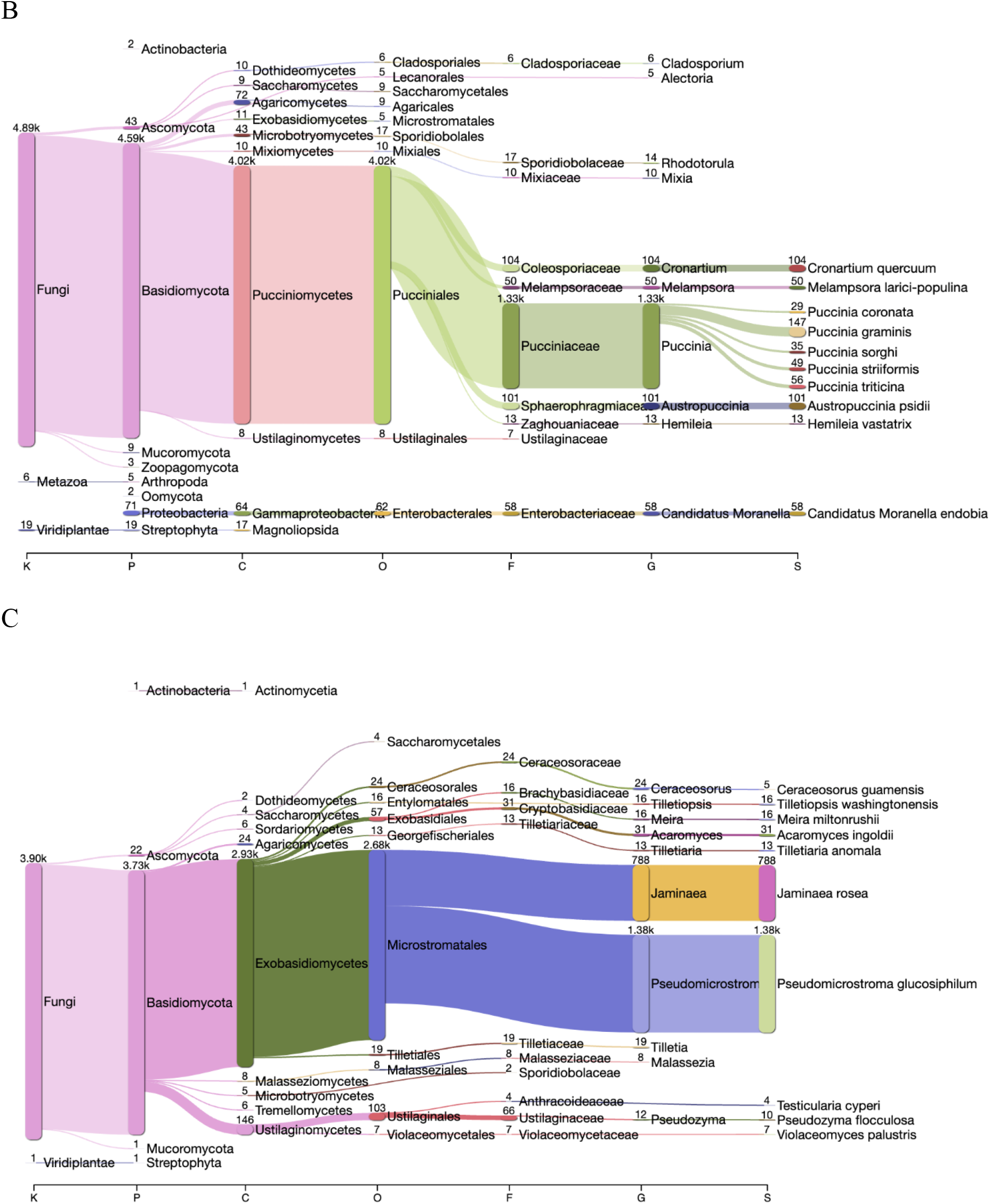
Taxonomic homology-based Taxolotl (v0.1.2) plots of the Hv178a genome with annotation from GeMoMa (v1.7.1) protein predictions for A) Complete sequence assembly, B) Hv178a genome and C) Hv178a associated fungal genome (AFG). Scale bar lettering represents Kingdom, Phyla. Class, Order, Family, Genus and species.

#### Read depth genome size prediction and copy number analysis

Genome size predictions were made with DepthSizer v1.6.3 (https://github.com/slimsuite/depthsizer), using partitioned PacBio reads (see below) mapped with Minimap v2.4 (Li 2018) and BUSCO v5 (Simão *et al*. 2015) gene predictions (see above). Read depth statistics and predicted copy numbers were generated using DepthKopy v1.0.3 (Chen *et al*. 2022).

#### Read partitioning

Following the identification of contig 002765F as putative mtDNA and SCAFF12 as a possible accessory genome, reads were partitioned into three subsets: Hv178a, Hv AFG, Hv178a mtDNA, and unmapped reads. PacBio long reads were mapped to the complete assembly using Minimap2 v2.22 (Li 2018). Illumina paired end reads were mapped using BWA-mem v0.7.17 (Li and Durbin 2010). Samtools v1.15 (Li *et al*. 2009) was then used to extract BAM files of aligned reads and fastq files of full-length mapped reads for each assembly subset.

#### Post-scaffolding polishing, contamination removal and cleanup

Genome polishing was attempted with Pilon (Walker *et al*. 2014), HyPo v1.0.3 (Breitwieser and Salzberg 2020) and Racon v1.4.5 (Vaser *et al*. 2017) and assessed using BUCO v5.3.0 (Simão *et al*. 2015) completeness. No polishing strategy was found to improve BUSCO Genome Completeness. Vector screening and removal of low-quality scaffolds was performed with Diploidocus v1.3.1 (Chen *et al*. 2022) using partitioned Hv178a reads and BUSCO results. Initial NCBI screening identified possible *Daucus carota* (wild carrot) contamination, restricted to a single contig of Scaffold 1. This contig was added to the Univec contamination database for Diploidocus contamination screening. Six contigs identified by Taxolotl as bacterial contaminants were removed prior to tidying. Diploidocus filtered 24 contigs from the genome as low quality, and a further 64 as probable haplotig false duplications.

#### Repeat family annotation

Transposable elements and simple sequence repeats were annotated with Repeat Modeler v2.0.1 (Smit and Hubley 2019) and Repeat Masker v4.1.0 (Smit *et al*. 2019). Initial repeat libraries were constructed for the complete draft assembly. Partitioned Hv178a and Hv178a AFG genomes were then annotated independently using the corresponding full haplotype repeat library.

#### rDNA prediction

Ribosomal rDNA genes were predicted with barrnap v0.9 (https://github.com/tseemann/barrnap (kingdom = eukaryote). Full-length rDNA repeats were identified by extracting annotations for 5.8S, 18S and 28S rRNA genes, excluding partial predictions, and identifying scaffolds containing multiple copies of all three. Full-length rDNA repeats were found on Hv178a scaffold 7, Hv178a AFG (scaffold 12), and an additional twenty contigs flagged as repetitive by Diploidocus. Predicted rRNA genes were searched against the NCBInr nt database with BLAST+ v2.11.0 blastn, confirming most contigs (and scaffold 7) as *Hemileia vastatrix*. Hv178a AFG returned high identity hits to the genus *Sympodiomycopsis* and *Malassezia*, giving possible insight into the taxonomic placement of this unknown yeast. Two contigs, CTG001507A and CTG001646A, had top-ranked hit to *Golovinomyces cichoracearum* mitochondrion (18S) and *Jaminaea angkoriensis* mitochondrion (28S). These contigs could represent unidentified Hv178a AFG mitochondrial sequence or may indicate that a low level of contamination may remain. One contig, CTG002427A, appears to contain rRNA repeats from a *Planococcus* species.

#### *Hemileia vastatrix* (CIFC Hv178a) scaffolded assembly statistics

The set of 11 ‘core’ scaffolds, plus an Hv178a associated fungal genome (Hv178a AFG, see below), as scaffold 12, were initially designated as the assembly and made available at 10.5281/zenodo.6757928 (v1.1). The Hi-C heatmap (Figure 2) showed that the Hv178a AFG scaffold has no interaction with the other scaffolds, indicating a likely second genome or subcellular compartment. Following further decontamination, the genome assembly, not incorporating the Hv178a AFG scaffold, was made available at 10.5281/zenodo.6816635 (v1.2) as detailed in the data availability section. The genome incorporates the fully circularised 171 kb mitochondrial DNA sequence as the final scaffold.

**Figure 2.**
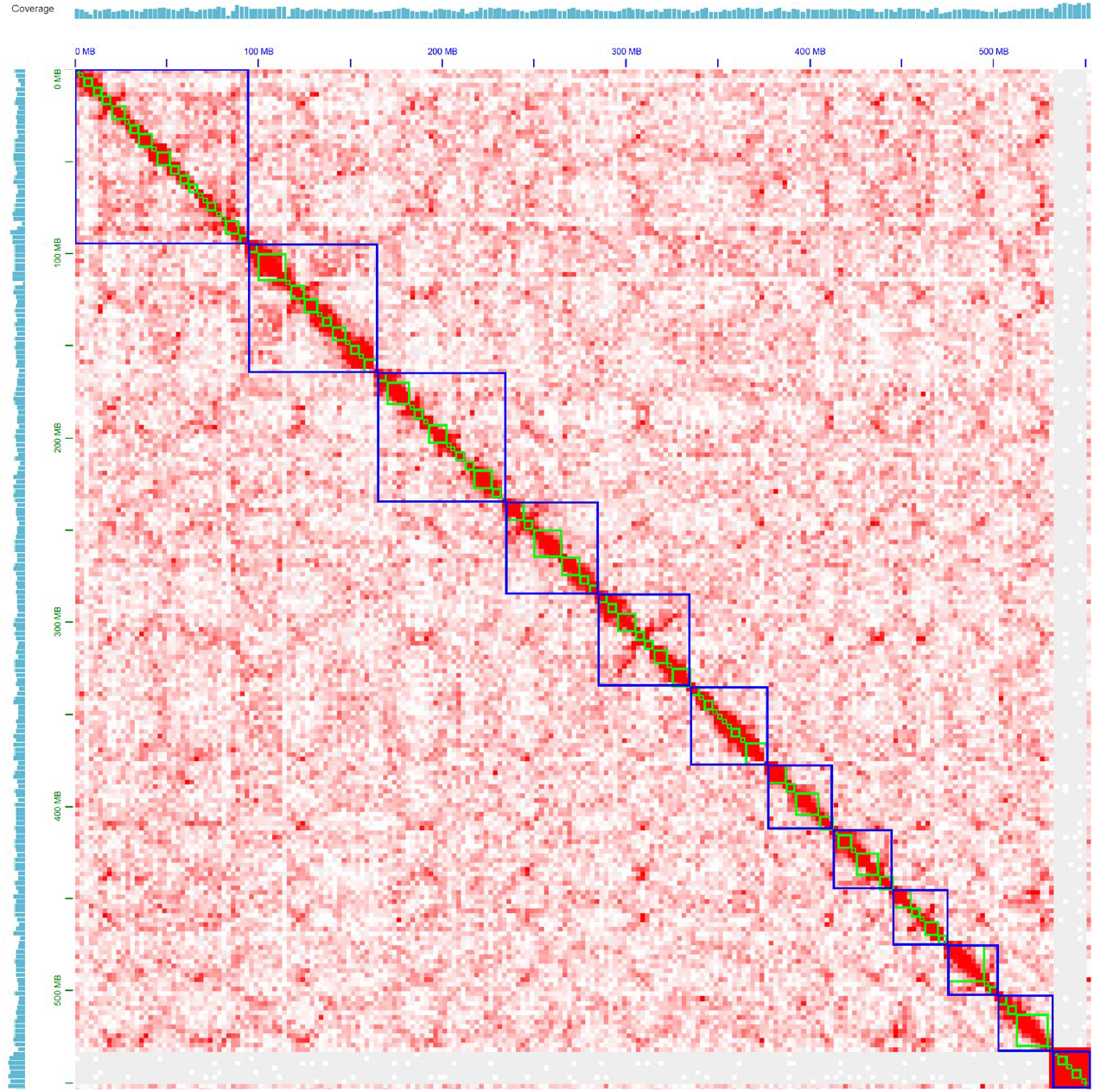
Image from Juicebox visualisation software (v 1.11.08) of the scaffolded *Hemileia vastatrix* (strain Hv178a) assembly heatmap shows 11 scaffolds and an additional 12^th^ scaffold that has no interaction with the other scaffolds, here termed Hv178a AFG (an associated fungal genome). Hi-C coverage is represented in blue.

**Table 1.** *Hemileia vastatrix* (strain Hv178a) Hi-C scaffolded genome assembly statistics using Quast (Gurevich *et al*. 2013) and BUSCO v. 5.1.2 (Simão *et al*. 2015) with basidiomycete_odb10 database. Hv178a plus AFG (an associated fungal genome) statistics are included as shaded cells.

| Genome | Contigs<br>(Scaffolds) | Total Length | Largest scaffold | N50 | L50 | GC (%) |
| --- | --- | --- | --- | --- | --- | --- |
| Hv178a | 57 (11) | 546,464,377 | 95,899,683 | 51,490,964 | 4 | 33.35 |
| Hv178a plus AFG | 58 (12) | 565,096,666 | 95,899,683 | 51,490,964 | 4 | 34.15 |
| BUSCO results | Complete | Single copy | Duplicated | Fragmented | Missing | Total(n) |
| Hv178a | 1,241<br>(70.4%) | 1,222<br>(69.3%) | 19<br>(1.1%) | 53<br>(3.0%) | 470<br>(26.6%) | 1,764 |
| Hv178a plus AFG | 1,692<br>(95.9%) | 799<br>(45.3%) | 893<br>(50.6%) | 13<br>(0.7%) | 59<br>(3.4%) | 1,764 |

## Discussion

### The *Hemileia vastatrix* (CIFC Hv178a) genome size conforms to Hv33

Our assembled genome represents 11 large pseudochromosomes with a total mean GC content of 33.35%. An additional 18 Mbp Hv178a associated fungal (AFG) scaffold was also assembled and is retained in our genome analyses. While our initial genome size estimate used the predicted 1C calculation determined with flow cytometry (FCM) of 796 Mb (Tavares *et al*. 2014), we consistently found our assemblies conforming to ∼ 540 Mbp, in accordance with Porto et al. (2019). Given that Tavares et al. (2014) also worked with the Hv178a isolate (http://www.zbi.ee/fungal-genomesize/?q#bobx) the size discrepancy is intriguing, but could relate to known errors that can occur using FCM for genome size estimation that are dependent on cell-cycle stage, the relative genome size for the host plant, and the nucleic acid stain (Wang *et al*. 2015; Ortega-Ortega *et al*. 2019). It is also possible that this is due to collapsing of highly repetitive regions during the assembly and scaffolding process. Statistics indicate that the Hv178a genome is 546 Mbp, when the scaffold containing the Hv178a AFG is excluded (Table 2).

### Hv178a chromosome numbers align with cytological studies

The final Hv178a assembly has 11 pseudochromosomes. *H. vastatrix* is an ancestral rust, from the family Mikronegeriaceae. Up to now, chromosome numbers have not been clearly assigned to this evolutionary group of rust-forming basidiomycetes. An earlier cytological study reported the difficulty in determining a conclusive number of chromosomes for *H. vastatrix* and suggested as the best resolved data the presence of 14 chromosomes (Chinnappa and Sreenivasan 1965). More recently, Tavares et al. (2013) determined between 7-13 chromosomes using both electrophoretic and cytological karyotyping, as well as fluorescence in-situ hybridization of two races of *H. vastatrix* (race II - isolate 1065 and race VI - isolate 71) from the CIFC/ISA collection. While several distantly related rust fungi within the Pucciniamycotina subdivision are known to have 18 haploid chromosomes (Schwessinger *et al*. 2020; Wu et al. 2021; Edwards *et al*. 2022), a much greater variation in chromosome number may be expected among the Pucciniales, particularly in ancient rusts.

### A Hv178a associated fungal genome (AFG) incorporates many conserved genes

The Hv178a AFG scaffold is an intriguing but unexplained finding from this assembly. We did not find this scaffold in a search within the genome of the Hv33 strain. While the taxonomic analysis and heatmap (Figures 1 and 2) and the BUSCO statistics (Table 3) suggest that this scaffold is a contaminant, we could not unambiguously attribute the sequence data to another organism. The scaffold shares taxonomic conservation with exobasidiomycetes and shares conserved BUSCO genes with basidiomycetes. Phylogenomic analysis with using BUSCO genes from Ensembl fungal genomes most closely associated Hv178a AFG with *Pseudomicrostroma glucosiphilum* (Kijpornyongpan and Aime 2017), a yeast (data not shown), whilst blastn analysis of predicted rRNA genes suggests it is a relative of *Sympodiomycopsis* or *Malassezia*. However, contamination was not detected with universal ITS primers during DNA preparation, nor was a second mitochondrial sequence observed within the total genome data, despite attempting to assemble using a seed read input from *P. glucosiphilum* cytochrome c oxidase (NCBI accession XM_025490016.1). Another interesting observation is that the removal of this scaffold led to incomplete conserved gene reporting for the Hv178a genome (Table 4), despite very high coverage of long read assembly data. Indeed, it appears that this scaffold was the source of the very high duplication rates hindering the initial deduplication with Purge Haplotigs software. We expect that further investigation is required to resolve the nature of this finding.

## Conclusion

We have assembled and curated the first scaffolded genome for the coffee leaf rust causing pathogen, *Hemileia vastatrix*. Our genome assembly for the strain CIFC Hv178a has shown that there are likely 11 haploid chromosomes. The genome resource, including a full mitochondrial sequence, will facilitate advances in understanding the pathogenicity of this virulent pathogen on *Coffea arabica*.

## Acknowledgements

Funding for the research was provided by PORLisboa, Portugal2020 and European Union through FEDER funds (LISBOA-01-0145-FEDER-029189) and by the Foundation for Science and Technology (FCT) through Portuguese funds (PTDC/ASP-PLA/29189/2017). We thank Ana Paula Pereira and the technical staff from CIFC/ISA, namely Célia Lopes, Idalina Gomes and Miguel Ribeiro, for the support provided on isolate multiplication and pathotype testing, as well as for the maintenance of coffee plants and preservation of CIFC rust collection.

## Author contributions

PAT assembled nuclear and mitochondrial genome assemblie, ran analyses, wrote much of the manuscript. RJE curated the genome data, ran analyses, prepared figures, contributed to the methods and results of the manuscript writing. PS ran genome assemblies. VI participated in the extraction and purification of HMW DNA. VV and MCS provided spore samples from CIFC’s collection, coordinated isolate multiplication and characterization, and contributed to the manuscript. HM scaffolded the genome. RFP supported the study and contributed to the manuscript. DB initiated and led the research, sourced grant funding, prepared samples for PacBio sequencing and for Hi-C and wrote much of the manuscript. All authors contributed to and read the manuscript.

## Data availability

Raw data is available at the following BioProject accession number at the National Centre for Biotechnology Information (NCBI): PRJNA837996. The complete genome, Hv178a_A.v1.2.fasta.gz, and associated fungal genome, Hv178a_AFG_A.v1.2.fasta.gz, are available here: 10.5281/zenodo.6816635. Note that the version **A.v1.2** is the current genome for this organism.

## Notes

https://zenodo.org/record/6816635#.YuSwKHZBxPY

https://www.ncbi.nlm.nih.gov/bioproject/PRJNA837996

### Competing Interest Statement

The authors have declared no competing interest.

